# Decoding Biomolecular Condensate Dynamics: An Energy Landscape Approach

**DOI:** 10.1101/2024.09.24.614805

**Authors:** Subhadip Biswas, Davit A Potoyan

**Affiliations:** Department of Chemistry, Iowa State University, Ames, IA 50011, USA; Department of Biochemistry, Biophysics and Molecular Biology, Iowa State University, Ames, IA, 50011, USA

## Abstract

A significant fraction of eukaryotic proteins contain low-complexity sequence elements with unknown functions. Many of these sequences are prone to form biomolecular condensates with unique material and dynamic properties. Mutations in low-complexity regions often result in abnormal phase transitions into pathological solid-like states. Therefore, understanding how the low-complexity sequence patterns encode the material properties of condensates is crucial for uncovering the cellular functions and evolutionary forces behind the emergence of low-complexity regions in proteins. In this work, we employ an alphabet-free energy landscape framework of the stickers and spacers to dissect how the low complexity patterns of proteins encode the material properties of condensates. We find a broad phase diagram of material properties determined by distinct energy landscape features, showing that periodic repeat motifs promote elastic-dominated while random sequences are viscous-dominated properties. We find that a certain degree of sticker periodicity is necessary to maintain the fluidity of condensates, preventing them from forming glassy or solid-like states. Finally, we show that the energy landscape framework captures viscoelastic trends seen in the recent experiments on prion domains and makes predictions for systematic variation of protein condensate viscoelasticity via altering the periodicity and strength of sticker motifs.

**TOC Graphic:** 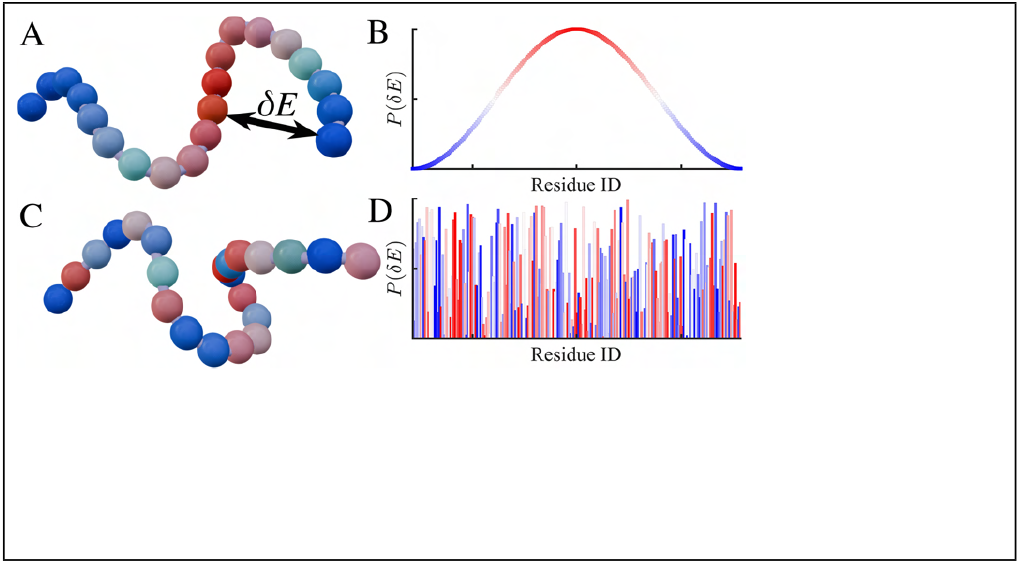

## Introduction

A significant fraction of the eukaryotic proteome contains sequences with quasi-periodic and low-complexity domains (LCDs).^1–3^ Proteins with LCDs display high conformational flexibility and, unsurprisingly, a high affinity for phase separation. Liquid-liquid phase separation (LLPS) of proteins has emerged as a ubiquitous mechanism for the biogenesis of biomolecular condensates and membranes organless with numerous organizational, regulatory, and signaling functions. ^1,4^ The material properties and structural organization of condensates appear tightly regulated in cells.^5,6^ Misregulation of these properties often leads to an irreversible transition to solid-like states associated with numerous pathological diseases.^7,8^ Additionally, material properties such as viscoelasticity, viscosity, and interfacial tension are sensitive to sequence mutations, particularly in regions containing low-complexity domains (LCDs).^9–15^ These observations have been rationalized by sticker and spacer models, which have provided numerous insights into sequence-condensate thermodynamic and dynamic properties of condensates.^16–18^ For instance, in recent reports, flow activation energies of various condensate phases were measured, finding that localized sticky motifs in sequences dramatically alter network reconfiguration times in condensates, thereby contributing to observed sequence dependence of viscosities.^10,19,20^ Nevertheless, the broader role of low-complexity sequence patterns in proteins and their connection with condensates’ material and structural features remains poorly understood. This is partly due to the absence of guiding evolutionary and biophysical principles for sequence grammar behind material properties. For foldable protein sequences, the energy landscape framework has provided a clear alphabet-free design principle;^21–23^ evolutionary pressure optimizes the energy gap between unfolded and folded states at physiologic conditions to avoid deep kinetic traps thereby ensuring rapid and robust folding. The key insight from energy landscape theory has revolutionized protein design and structure prediction in the last decade.^24–26^ It is therefore intriguing to search for a similar conceptual and computational toolbox couched in the language of energy landscapes ^27–30^ which can help in identifying design principles of condensates with material properties compatible with functional dynamics.

Such principles would help target the formation of aberrant phases with solid-like material properties implicated in several neurodegenerative diseases and the design of novel condensates. To dissect the role of low-complexity periodic repeat motifs on condensates’ thermodynamic and dynamic characteristics, here we introduce an Alphabet-Free Energy Landscape framework of Stickers and Spacers (AFELAS) (Fig 1). In this alphabet-free energy landscape model, protein sequence is defined by specifying the relative “sticky” affinity of residues with respect to baseline “spacer” affinity *δE* = *ε*_*i*_ − *ε*_*sp*_. Armed with an energy landscape framework, we dissect the contribution of sequence complexity and periodicity of sticker repeat domains (Fig 1A-D) in dynamic and viscoelastic properties and shed light on recent experiments probing sequence-dependent material properties of condensates. ^10,31^

**Figure 1:**
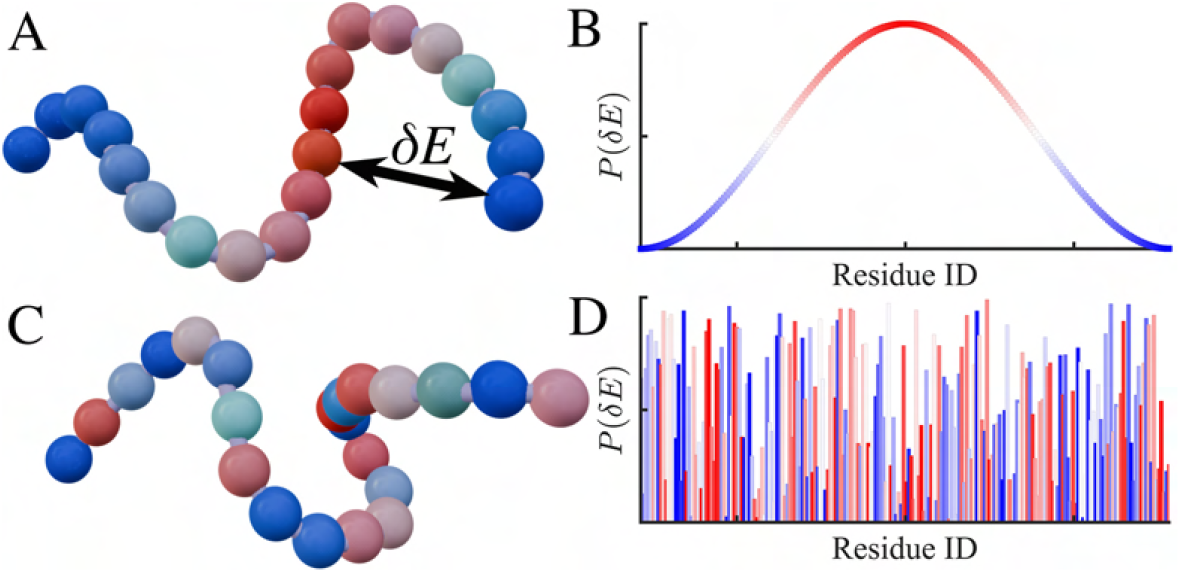
(A) Schematic depiction of an alphabet-free sticker spacer framework of protein chains where associative interactions between domains are sampled from an energy landscape. The blue-red color scheme represents distinct motif IDs, where blue indicates spacers and shaded blue-red represents stickers with varying interaction energies. (B) The extreme case of a periodic arrangement of stickers corresponding to chain (A) where the probability of non-bonded interaction energy is distributed with a single unique period along the chain (C) The extreme case of randomly distributed interaction energies along the chain (D).

We discover that a certain degree of sticker periodicity is necessary to maintain the fluidity of condensates, preventing them from forming glassy or solid-like states. Through simulations employing an energy landscape framework of stickers and spacers, we also shed light on the mechanisms of condensate formation, revealing a progression from fluid-like network conformations to viscoelastic systems in the dilute and dense phases, irrespective of the protein sequence.

### Alphabet-Free Energy Landscape of Stickers and Spacers (AFELAS)

Here, we describe the physical motivational and computational details of the energy landscape-based sticker and spacer model (Fig 1). The key idea is adopting a relative energy scale that differentiates between sticker and spacer regions, which captures continuous sequence variation without operating with an explicit alphabet of protein residues. The protein chains are coarse-grained into beads connected by anharmonic FENE (Finite Extensible Nonlinear Elastic) springs, described by the potential 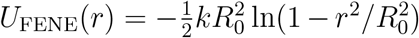. Here, *r* represents the distance between adjacent beads, with a spring coefficient of *k* = 20*ϵ/σ*^2^ and maximum extensibility of *R*_0_ = 1.5*σ*, where *σ* denotes the bead diameter, establishing the length scale for LJ (Lennard-Jones) simulations. Bond angles between consecutive beads are constrained by an angular harmonic potential, *U*_*θ*_ = *K*_*θ*_(*θ* − *θ*_0_)^2^, where *K*_*θ*_ denotes the potential energy and *θ*_0_ = 150^*°*^ is the equilibrium angle. Non-bonded interactions are incorporated through the Lennard-Jones (LJ) potential, expressed as 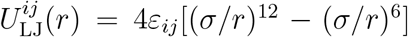, effective within a cutoff distance *r*_c_, where *U*_LJ_(*r*) linearly goes to zero at *r*_*c*_.

The strengths of non-bonded interactions *ε*_*ij*_ are calibrated to range between well depth 0 (WCA) and *kT*, following a periodic modulation *ε*_*ii*_ = sin^2^(2*π*Res_*i*_*/k*), Res_*i*_ with period *k* set at 10, 25 and 100 in Fig. (2 A, B). Inter-motif interactions *i* ≠ *j*, where the energy between *i* and *j* monomers is weaker than intra-motif interactions (*i* = *j*), are established via *ε*_*ij*_ = *ε*_*ii*_*ε*_*jj*_, as illustrated in the pairwise interaction energy landscape diagram in Fig. (2 C).

The interaction energies with the well depths *ϵ*_*ij*_ ≤ 0.1 are set to 0, representing WCA repulsive interactions, and are termed “spacers”, while the beads with attractive interactions are referred to as “stickers”. The functional form ensures that the total energy across different periodicities remains the same, allowing one to dissect the impact of sequence patterns on the viscoelastic and structural properties of condensates. We have considered the following sticker spacer motifs using a notation of *P*_*k*_ denoting sequence energy landscape with sticker periodicity *k* = 10, 25, 100 (Fig 2 A). We have then introduced randomness in sequence in steps, denoting 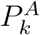 where sticker motifs are randomized while fixing spacers, 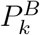 where alternatively higher interaction strength sticker - spacers or lower interaction strength sticker motifs are distributed along the chain (Fig. 4). Finally, taking the very same periodic chains and randomly arranged along the chain backbone where the extreme random heteropolymers denoted as *R*_*k*_ (Fig 2 A). Simulation snapshots illustrating these configurations of the bulk solution in equilibrium are presented in Fig. 2 D.

**Figure 2:**
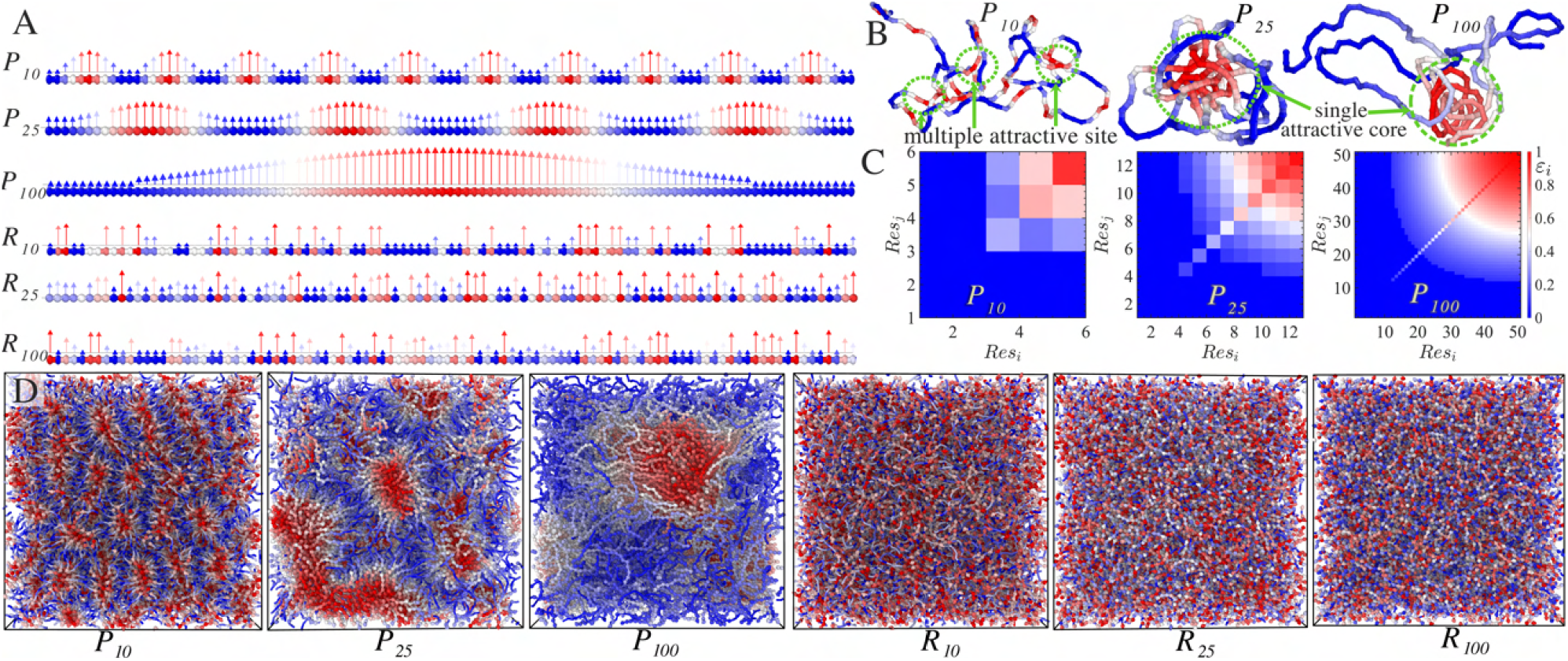
Schematic description and simulation snapshots of the sequence-specific spatial arrangement of stickers and spacers within the chains. The variations of i) periodically distributed sticker-spacer motifs *P*_10_, *P*_25_, and *P*_100_, and ii) randomly distributed sticker-spacer motifs along the chain backbone *R*_10_, *R*_25_, and *R*_100_ are shown in (A). Conformations of the AFELAS chain in very dilute solutions are depicted in (B). The interaction strengths between distinct stickers are illustrated in the energy landscape diagram. The pairwise interaction matrix in (C) depicts the red motif exhibiting an interaction energy of *ε* = 1, representing the highest sticker strengths, while a blue motif denotes spacers with the lowest interaction energy. (D) As the periodicity of the sticky region along the chain increases, intermediate sticker strengths are introduced, conserving total energy along the chain. Simulation snapshots of the bulk system of AFELAS chains with the aforementioned periodic and randomly distributed stickers while fixing all other parameters. A membrane-shaped layer forms for *P*_10_, a cluster domain for *P*_25_, and a larger sticky cluster for *P*_100_. The structural architecture transitions into a homogeneous condensed phase for randomly distributed *R*_10_, *P*_25_, and *R*_100_.

## Results

### Thermodynamics and Dynamics of low complexity periodic motifs vs random heteropolymers

Before delving into material properties encoded by energy landscapes of stickers and spacers, we first analyze how patterns of stickers and spacers encode the thermodynamics of phase equilibrium. All the simulated chains have the same length *N*_*p*_ = 200, mimicking typical condensate-forming proteins with multiple LCDs. We systematically vary the periodicity of the sticker motifs (where Res_Period_ = 10, 25, 100) while keeping the total “sticker energy” conserved. Despite having constant sticker energy, we find that the critical temperatures exhibit significant variation with sticker periodicity (Fig. 3 A). For shorter period sequences, *i*.*e*., *P*_10_, the critical temperature is *T*_*c*_ ≈ 1.4*ϵ*. For longer periodic sequences like *P*_25_, the critical temperature increases to *T*_*c*_ ≈ 2*ϵ*. In the case of much longer period *P*_100_, *T*_*c*_ is almost twice as high as that of the *P*_10_ sequence arrangements, *T*_*c*_ ≈ 2.7*ϵ* (Fig. 3 A).

**Figure 3:**
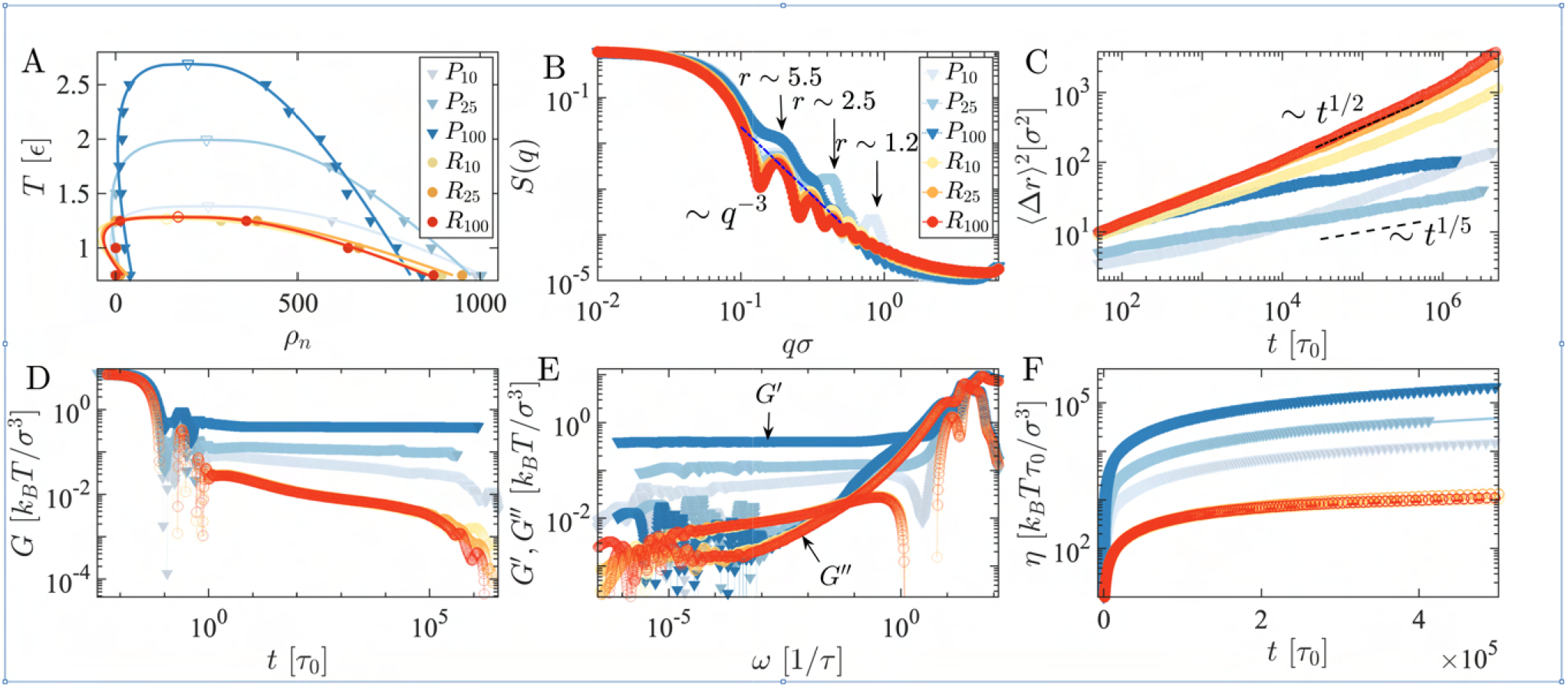
Equilibrium and non-equilibrium properties of the programmable energy landscape sticker-spacer model: A) The phase diagram of periodically distributed chains (blue shaded) condensates reveals that *P*_100_ exhibits a critical temperature *T*_*c*_ more than two-fold higher than that of *P*_10_ due to the lower periodic occurrence of the sticker. Conversely, the randomly distributed (red shaded) for the same systems shows a lower critical temperature *T*_*c*_. B) Anomalous peaks in the structure factor *S*(*q*) of periodically distributed chains *P*_period_ characterize equilibrium folded sticky domain sizes absent in randomly distributed chains. C) With increasing period, periodically distributed chains *P*_period_ displays more glassy arrested behavior, following *t*^1*/*5^, whereas randomly distributed chains follows *t*^1*/*2^. D) The complex modulus *G* of randomly distributed chains decays faster than periodically distributed chains. E) Elastic (*G*^*′*^) and viscous (*G*^*′′*^) moduli exhibit a high-frequency viscous-dominated regime and an elastic-dominated crossover at an intermediate frequency range. For *ω* → 0, a Maxwell viscous-dominated trend is observed for randomly distributed chains, while a Kelvin-Voigt *G*^*′*^ plateau is observed for periodically distributed chains. F) periodically distributed systems are highly viscous, with viscosity increasing with the periodicity of the stickers, whereas randomly distributed systems have an order of magnitude lower viscosity.

**Figure 4:**
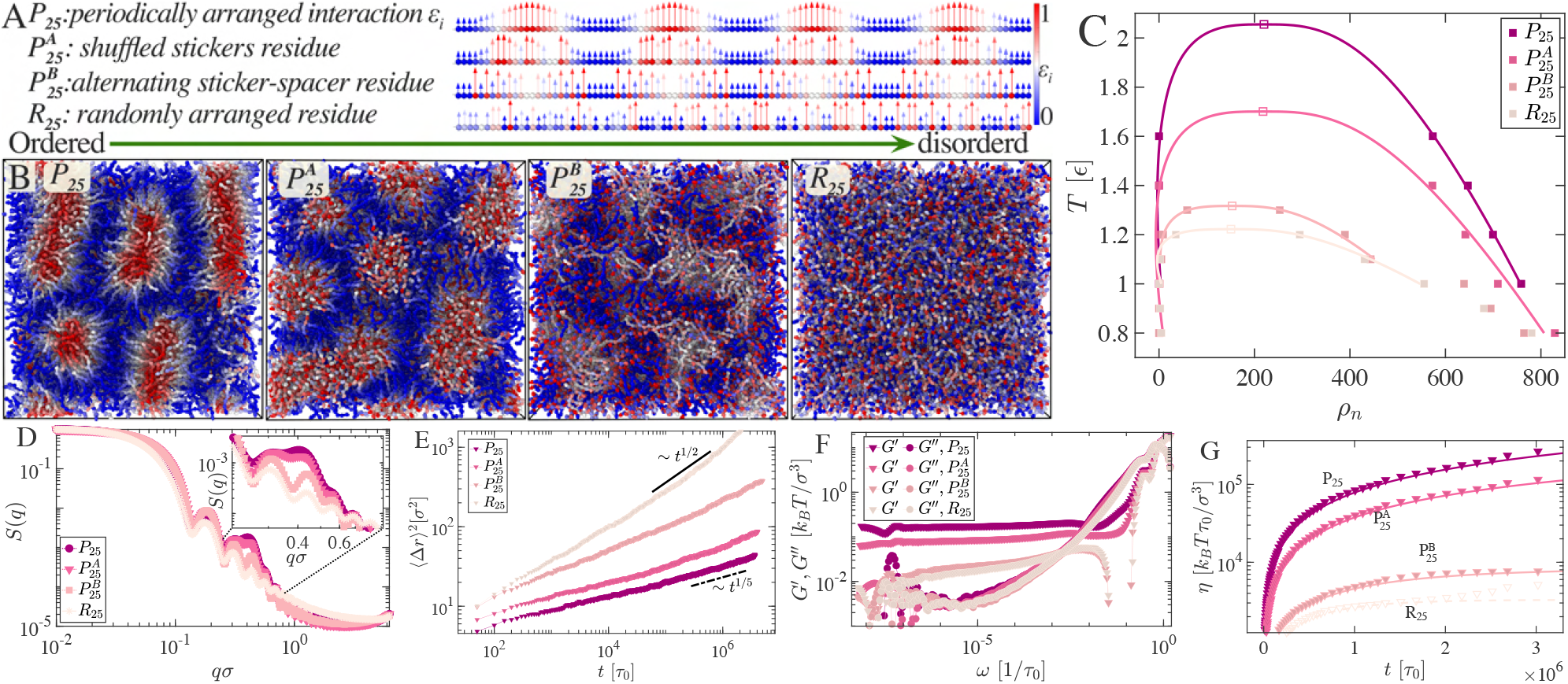
Simulation snapshots depict the transition from ordered to disordered energy landscapes of monomer interactions for various sticker and spacer arrangements. Alongside periodically distributed *P*_25_ (A & B) and randomly distributed *R*_25_ (A & B) chains, two new sticker-spacer arrangements are introduced. 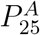 (A): sticker residues are shuffled while keeping spacer periodicity fixed, and its corresponding bulk simulation snapshot is shown in (B). 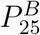 (A & B): an alternative sticker-spacer chain arrangement and its corresponding bulk simulation snapshot shown here. (C) shows the phase diagram of the four different sequences pattering where *T*_*c*_ of *P*_25_ is almost double of *R*_25_. (D) Static structure factors *S*(*q*) for condensates formed by different sticker-spacer arrangements. (E) Mean square displacement averaged over all beads and shown for condensates formed by different sticker-spacer arrangements. (F) Elastic and viscous moduli for different sticker-spacer arrangements. (G) The viscosity was computed for the different sticker-spacer arrangements.

As the extreme case, we examine the fully randomized arrangement of sticker and spacer motifs denoted as *R*_10_, *R*_25_, &*R*_100_. Intriguingly, regardless of the permutation of the periodically distributed chain, the phase diagram of the randomly arranged sequences manifests approximately the same critical temperatures, denoted as *T*_*c*_ ≈ 1.4*ϵ*, (Fig. 3 A). Notably, a mere shuffling of the sticker and spacer motifs of *P*_100_ results in a halving of the critical temperature from *T*_*c*_ = 2.7*ϵ* to *T*_*c*_ = 1.4*ϵ*. Observing the phase diagram, one notices that periodically arranged stickers demonstrate a greater propensity for phase separation than randomly distributed stickers. Moreover, phases will likely be more stable at higher temperatures if the periodicity broadens. Augmenting the periodicity of attractive beads within our model resembles increasing the number of stickers, akin to folded binding domains, in a protein structure. ^16,32^ Experiments have demonstrated that mutations at sticker sites, rendering them functionally inert, reduce the tendency of proteins to undergo phase separation,^33^ a phenomenon analogous to our findings.

### Structure factor

Here, we explore the correlation between sticker spacer arrangements and the emergence of local structural order. We observe an increase in the thickness of the sticker layer or domain size with elongated periodic chains, resulting in a shift of scattering peaks towards lower *q*-values (see in Fig. (3 B)). In randomly distributed chains, *S*(*q*) decays as *q*^−3^, without anomalies in the peak, while in periodically distributed chains, one peak is significantly higher, indicating local structures of sticker domains. The reciprocal of the peak position in Fourier space corresponds to the thickness layer (*r* ≈ 1.2*σ*) in *P*_10_, which increases longer periodicity for *P*_25_ (*r* ≈ 2.5*σ*), and further increases for *P*_100_, the shift to even lower values (*r* ≈ 5*σ*), as indicated in Fig. (3 B). The *S*(*q*) reveals the condensates’ local architecture, which, as we shall see in subsequent sections, influences nonequilibrium rheological and viscoelastic properties. An increase in sticky domain size suggests the creation of folded sticky domains, enhancing stability and promoting a more viscoelastic behavior. Conversely, randomly distributed chains exhibit fluidic behavior due to the absence of local structure in the condensed phase. Equilibrium contact map calculations of condensates support the local structures seen in *S*(*q*) peaks. Specifically, the contact maps illustrate spatial proximity (*r*_*c*_ *<* 2*σ*) between motifs *i* and *j* within a biomolecular condensate (Fig S1). In periodically distributed chains, well-formed domains of sticker motif contacts are evident, while consistent contact between spacers is lacking. Conversely, randomly distributed chains exhibit no consistent contact regions capable of forming sticker domains.

### Mean Squared Displacement

We use mean squared displacement (MSD) to elucidate the arrested subdiffusive nature of condensates from different sticker spacer patterning. The MSD, ⟨Δ*r*^2^(*t*)⟩ = *Dt*^*α*^, where *D* is related to the diffusion constant and *α* is the scaling exponent. We observe subdiffusive behavior, characterized by MSD scaling as Δ*r*^2^ ∼ *t*^1*/*2^ to *t*^1*/*5^, which arises from sticky domain formation and caging dynamics (Fig. 3 C). For randomly organized sticker spacer complex systems, the MSD exhibits a *t*^1*/*2^ scaling reflecting slowed molecular motion due to topological entanglements and localized interactions. A recent study has shown subdiffusion within protein droplets due to their viscoelastic nature, along with the significant difference in protein diffusivity between the interior and interface of the droplet.^34^ This phenomenon is consistent with a different periodicity of the sticker molecules, while arrangements of the stickers are randomly distributed along the chain in Fig. (3 C). As the degree of periodicity broadens, sticky clusters grow, and spatiotemporally arrested condensed phases exhibit slower dynamics (see blue lines in Fig. 3 C).

### Viscoelasticity

Recent microrheology experiments have demonstrated that the material properties of biomolecular condensates can vary widely, ranging from intricate Maxwell fluids to glassy or Kelvin-Voigt viscoelastic gels, depending on their composition and maturation times.^19,35–37^ Here, we elucidate how the energy landscape of sticker spacers encodes the diverse viscoelastic properties observed in experiments. We find that randomly distributed stickers (Fig. 3 D) show multiple relaxation times seen by fitting the complex modulus *G* to the generalized Maxwell model *G* = Σ_*i*_ *G*_*i*_ exp (−*t/τ*_*i*_). The complex modulus *G* goes to zero for all three randomly distributed sticker cases at long *t*, revealing Maxwell fluid behavior. Maxwell-like behavior persists also for sticker-spacer systems with short periodicity *P*_10_ (Fig. 3 D). However, broadening the period of sticker domains (*P*_25_ & *P*_100_) changes the material properties of condensate from Maxwell fluid to Kelvin-Voigt elastic solid behavior. The complex modulus for *P*_25_ & *P*_100_ gets nearly saturated at long time scales (Fig. 3 D).

The Fourier transform of *P*_25_ & *P*_100_ shows that the elastic modulus *G*^*′*^ is independent of imposed deformation frequency *ω* at long times (Fig. 3 E) a signature of Kelvin-Voigt solid nature of the condensed phase. As sticker domain size increases, entangled sticker segments fold back to make a strongly elastic-dominated viscoelastic condensed phase, where *G*^*′*^ of *P*_100_ is an order of magnitude larger relative to its randomly distributed *R*_100_ sequence. Note that, even though the elastic dominance differs for different arrangements of the sticker and spacers, viscous response, *G*^*′′*^, collapses on the same curve for all systems. The scaling behavior of Maxwell fluid in the viscous dominated long time scale regime indicates that as *ω* → 0, exhibits *G*^*′*^ ∼ *ω*^2^ and *G*^*′′*^ ∼ *ω*. Thus, one can think of the aging of biomolecular condensates as rearranging the sticker residue for a long time, which makes the systems fluidic to an elastic-dominated Kelvin-Voigt nature. ^18^ Our simulation reveals a strong correlation between the sticker profile and bulk viscosity of condensates (Fig. 3 F), linked across scales using Rouse theory.^38^ The periodic arrangement of the stickers results in an order of magnitude more viscous (Fig. 3 F) phase compared to their random counterparts. This is consistent with its thermodynamically more stable phases and structurally larger sticky domains formed by chains with periodically arranged stickers.

### Dynamical impact of local randomness in periodic stickers and spacers motifs

Having discussed the two extreme limits of periodic and random sticker patterns, we now turn to examine two intermediate cases of the disorder in stickers and spacers denoted as *P*_*A*_ and *P*_*B*_(Fig. 4 A & B) respectively. We use the 25-period residue sequence as the reference for comparison. In the case of 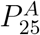, we fix the spacers defined by the lowest *ε* interaction strengths and randomize sticky domains defined by the higher values of *ε*. In case 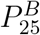, the sticker period is fixed, but the strengths of residues in the sticker domain alternate between higher and lower values sequentially (Fig. 4 A). This corresponds to typical *RGRG, Y GY G* motifs in many phase-separating proteins.^1,39,40^

Simulations show that one can have very distinct bulk equilibrium structures and material properties by simply tuning the degree of order in sticker and spacer domains (Fig. 4 B). Chains with periodically arranged ordered sticky regions self-assembled make micell-like architecture where the bulk is heterogeneously distributed with the sticker types (Fig. 4 A & B). A disordered sticker arrangement makes a structure-less, unstable, homogeneous melt (Fig. 4 B). In the intermediate states, 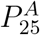&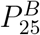, there are also structures present; however, they are not as prominent as *P*_25_ Fig. (4 A & B).

The phase diagram of these different sticker arrangements (Fig. 4 C) shows that extreme ordered *P*_25_ and random arrangement of stickers possess the most stable and least stable phases, respectively. The 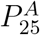&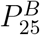 sequences lie in the intermediate stability regime. To structurally distinguish the architecture of these condensates, we calculate the structure factor *S*(*q*). An anomalous peak in the case of 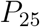&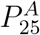 indicates the existence of local structure in the condensed phase. The peak position at *q* refers to the size of the sticker micell-like local structures (Fig. 4 D).

The prominent peak is lowered in the case of 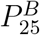, suggesting the local structure is fading out. In the case of a fully randomized sequence *R*_25_, one ends up with a homogeneously distributed melt system. The emergence of structural order is also reflected in subdiffusive dynamics. The exponent of the MSD decays with periodically arranged stickers ⟨Δ*r*^2^⟩ ∼ *t*^1*/*5^ (Fig. 4 E).

In terms of viscoelastic properties quantified by *G*^*′*^, *G*^*′′*^ (Fig. 4 F), we find that local structural order leads to elevated elastic modulus (Fig. 4 F). We plotted viscosity over time, whereas the saturation value as *G*^*^ → 0, *η* → *η*_0_ describes the total viscosity of the system. In condensates with periodically arranged stickers, *P*_25_ condensates form the most viscous systems, whereas it is lowest for *R*_25_ (Fig. 4 G).

One can also look into the contact time autocorrelation function, *C*(*t*), which tracks the dynamics of contact formation and offers insights into the viscoelasticity.^41^ Short timescales show rapid decay in *C*(*t*), suggesting fluid-like behavior with frequent contact disruptions and reforms. On longer timescales, *C*(*t*) indicates a shift toward a gel-like arrested state, where sustained interactions enhance viscosity and structural rigidity, mainly due to sticker-sticker interactions (see SI, Fig S4). Periodic sticker arrangements result in longer correlation times in *C*(*t*), while higher temperatures accelerate correlation decay, indicating reduced viscoelastic stabilization from sticker interactions (see SI Fig. S4).

### Comparison with experiments

We have reported that viscoelasticity grows as a function of the periodicity of stickers and that mutations in sticker regions have the most dramatic impact on material properties. In this section, we show that these results fall within the expected range of experimentally characterized material properties despite the limited data available in the literature on the viscoelastic properties of protein condensates with varying sticker and spacer residues. ^19,42^

We define periodicity for a real protein sequence to facilitate comparison with our energy landscape predictions. For this, we use a correlation length scale where the maximum energy strengths as defined by hydropathy index^43^ *λ*_*max*_, decays to half-width total maxima along the chain, ⟨*κ*⟩. We plot the inverse loss tangent, 1*/* tan *δ*, or the ratio of *G*^*′*^*/G*^*′′*^, as a function of ⟨*κ*⟩ Σ *λ*_*i*_*/λ*_*max*_*/N*_*p*_, where the chain length is *N*_*p*_ and *λ*_*i*_ is the energy amplitude of the *i*-th residue, and *λ*_*max*_ is the maximum energy among the sequence residues. Note that most of the experiments have been conducted with changing residues; therefore, along with the periodicity ⟨*κ*⟩, we also incorporate the unitless interaction energy parameter *λ*. A similar increasing trend in the elastic modulus with increasing periodicity is observed in our simulation data (Fig. 5 B). It is consistent with the limited experimental data from ^19^ (Fig. 5 A).

**Figure 5:**
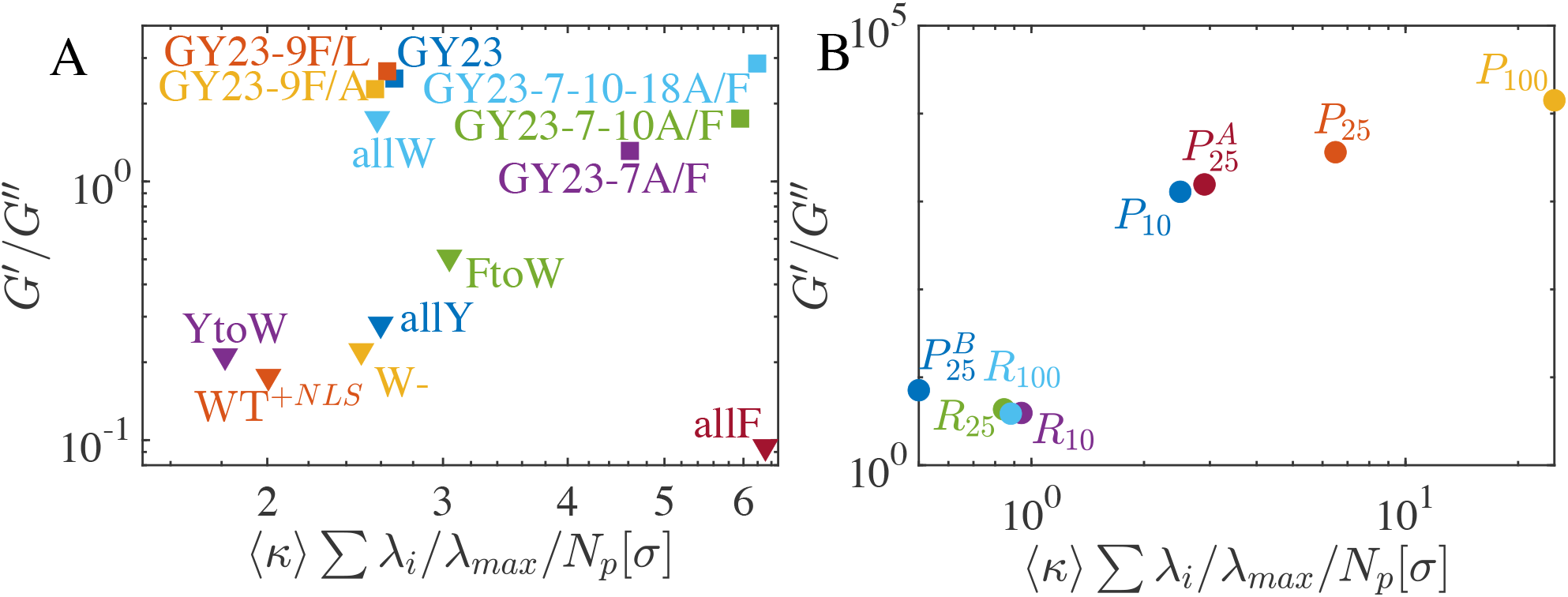
Comparison of the experimental results of *G*^*′*^*/G*^*′′*^ reported in^19^ and^42^ in (A) with our model of periodic and random sequences in (B) as a function of periodicity of the sequence. Both experimental and simulation results show an increase in viscoelasticity as the periodicity of the arranged sequences increases.

### Orientational order and correlations

A semi-flexible homopolymer chain’s persistence length (𝓁_*p*_) exhibits resistance to bending, which characterizes the mechanical and viscoelastic properties of constitutive bulk systems. The angle between a tangent vector 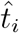 at position *i* and another tangent vector 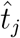 at a distance along the polymer contour, *s*, determines the correlation 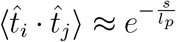, indicating the extent (*e*.*g*., persistence length, 𝓁_*p*_) over which correlations along the contour approach zero. Typically, the homopolymer with bending rigidity (or persistence length *L >* 𝓁_*p*_ *> σ*) reveals viscoelastic properties. As a reference, we note that tangent correlation (exponential decay) along the chain contour is spatially invariant in the case of homopolymer, as every bead is identical in nature.

In our model, even though the chain is not homopolymeric, we find a single exponential decay tangent correlation for random sequences as depicted in Fig. (6 D-F). Interestingly, sticky residues exhibit different correlation characteristics in the case of periodic sticker-spacer motifs. Due to the formation of clusters among sticky residues, there is strong anti-correlation behavior. ^44–46^ The orientational ordering in periodically arranged sticker systems reveals that certain beads show non-monotonous decay in tangent correlation. For spacer beads with repulsive interactions, tangent correlation decays exponentially. One sees long-distance ordering as a function of sticker interaction strength, which also leads to negative correlations, dictating that chains are folding back. The strength of interaction among sticker residues does not solely dictate the arrangement of folded sections (Fig. 6). Still, it is also influenced by the spatial distribution of the sticker beads (Fig. 6 B, C).

**Figure 6:**
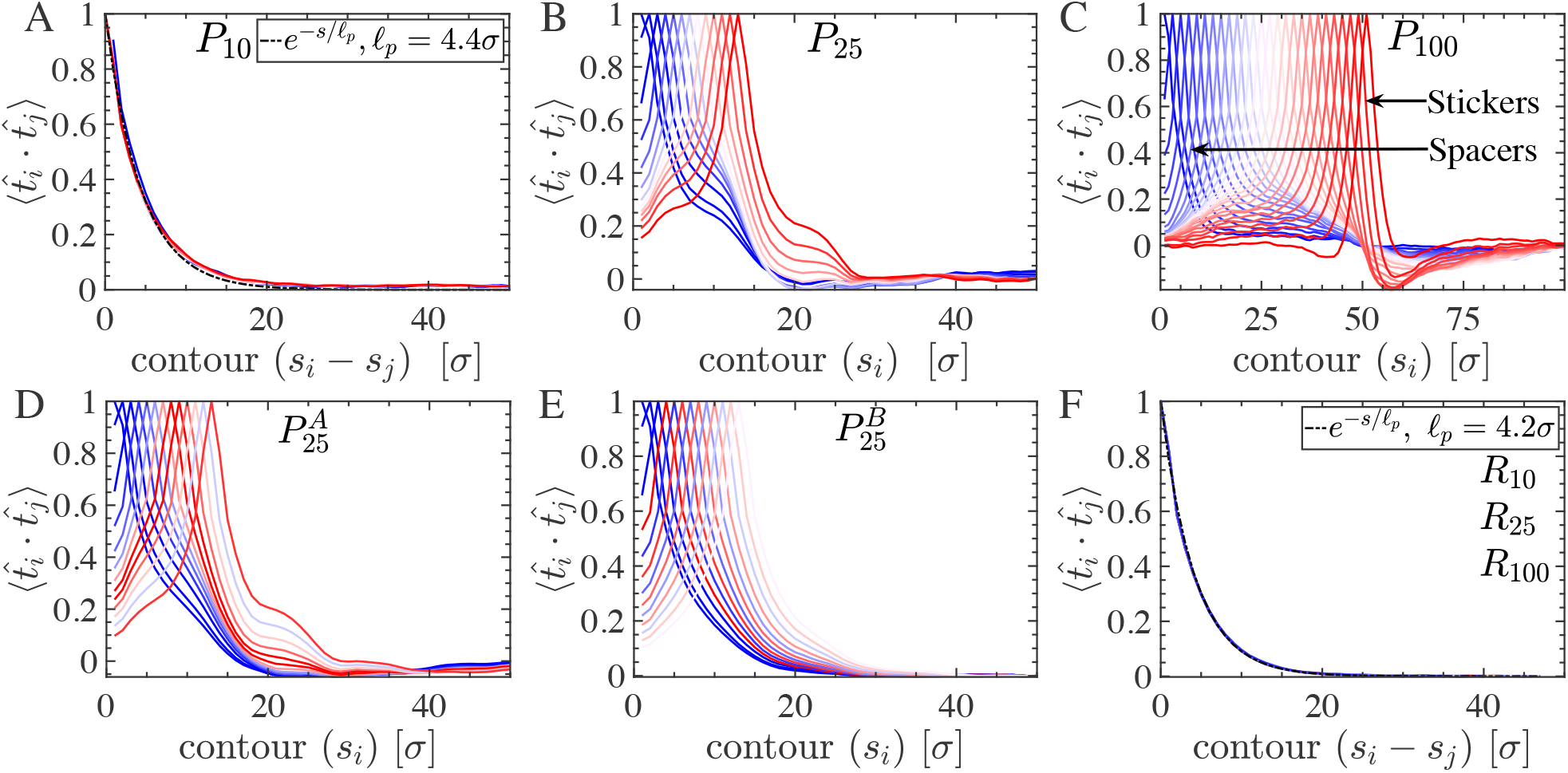
Tangent-tangent correlation in condensates was calculated for the periodically distributed (A-E) and randomly distributed (F) stickers. (A) For *P*_10_, periodic sticky regions aggregate; however, the chains show uniform exponential decay correlation. (B) *P*_25_ shows anomalous exponential decay in 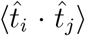, indicating a folded structure. (C) *P*_100_ displays diverse position-dependent residue characteristics and long-range order; spacer residues show exponential decay, while spool-like folded domains show non-exponential decay and negative correlation. (D) 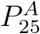 also shows anomalous exponential decay, and (E) 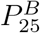 has distinct exponential decay lengths. (F) Randomly distributed chains (*R*_10_, *R*_25_, *R*_100_) show worm-like chain behavior with a persistence length 𝓁_*p*_ = 4.2*σ*.

The tangent correlation for short periodic sticker (*P*_10_) systems exponentially decays, revealing non folded sticky patch region (Fig. 6 A). Therefore, these systems do not form tightly connected sticky domains and exhibit viscous-dominated rheological behavior under applied deformation. Consequently, they are also less stable at high temperatures, leading to a lower critical temperature (*T*_*c*_) in the phase diagram than systems with broadly distributed periodic sticker beads. Systematic increase in periodicity of the sticker in Fig. (6 B & C), sticker residues with 0.5*ϵ*_*ij*_ *<* 1 have a higher tendency to fold back compared to spacers *ϵ*_*ij*_ ≈ 0, and those have *ϵ*_*ij*_ ≈ 1. The radius of curvature of the coiled folded sticky regions can be extracted from the minimum correlation distance. Spacer residues exhibit the expected exponential decay characteristics (Fig. 6 C); however, with an incremental increase in sticker strength, residues tend to accumulate within a sticky domain in a folded configuration.

Periodically arranged sticker molecules show arrested dynamics, whereas the randomly distributed stickers are more fluid in nature. We calculate the self-van Hove function to differentiate the spatial-temporal characteristics that resemble the distinct sub-diffusive nature (different exponent) of the dynamics. The self-part of the van Hove function 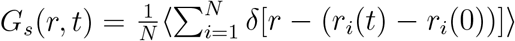 (where *N* is the total number of beads, *δ* is Dirac delta and ensemble average ⟨…⟩ over multiple trajectories), captures the probability distribution of particle displacements over a time interval *t*.^47^ The approximation of the self-van Hove function is Gaussian 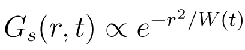.^48^ Unlike the Gaussian *G*_*s*_ (*r, t*) observed in structurally simple fluids,^48^ our analysis reveals nuanced characteristics, particularly in randomly distributed frustrated sticker-spacer condensed phase, where exponential decay tails signify anomalous behavior.^46,49,50^ In the case of periodically arranged systems, the self-van Hove function is Gaussian in nature, in early (dashed line Gaussian fit) and as late as *t* ∼ 10^8^*τ* (Fig. 7 A, B & C). On the other hand, randomly arranged stickers remain Gaussian in nature (Fig. 7 D, E & F). Therefore, we can conclude sticker beads are localized in the case of periodic stickers, as seen in (Fig. 7 A, B, C) compared to the randomly placed sticker-spacer. This temporal evolution of *G*(*r, t*) unveils invaluable information regarding the evolving microenvironment surrounding each sticky core, shedding light on structural rearrangements within the cluster. ^47^

**Figure 7:**
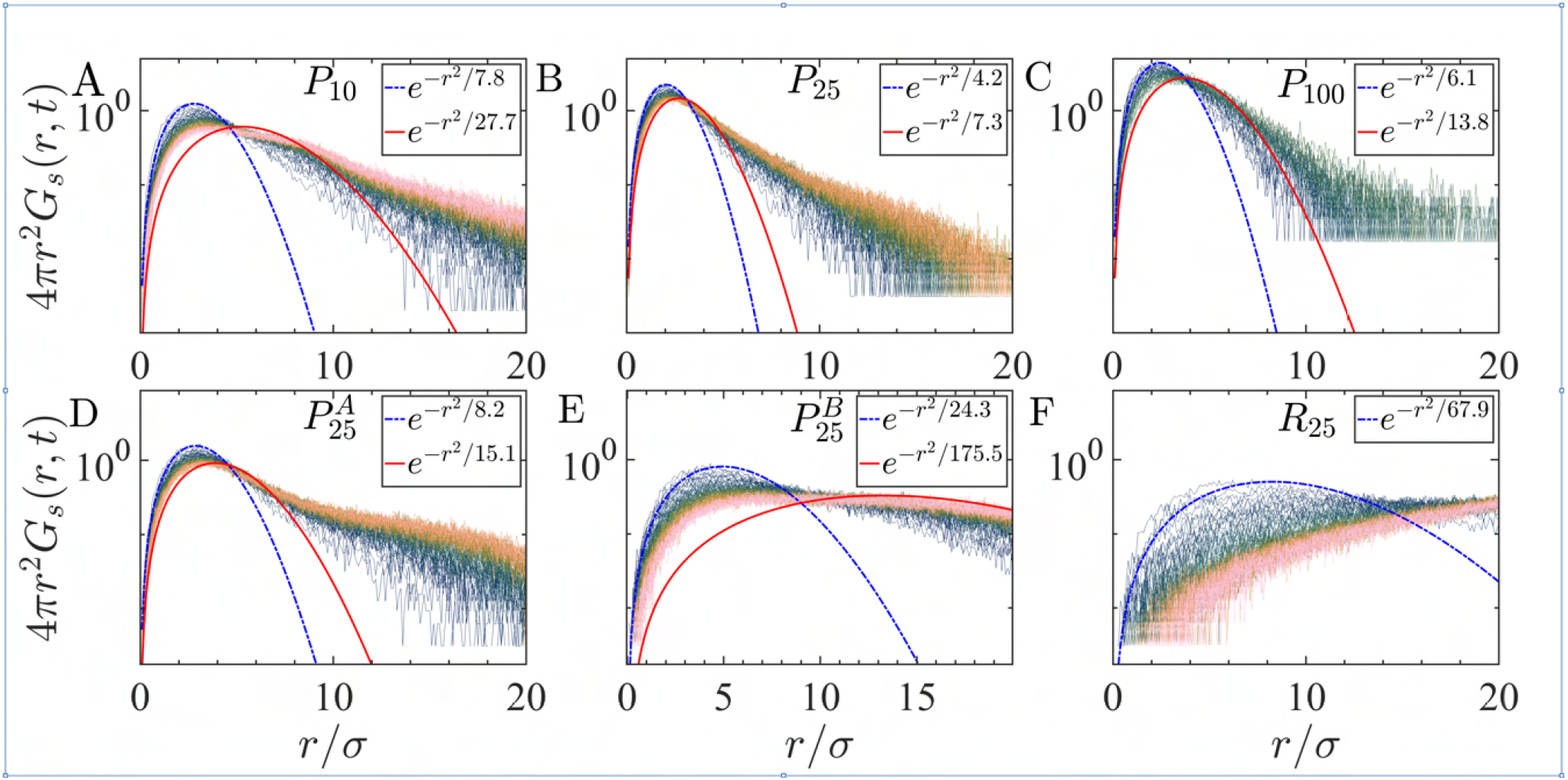
The self part of the van Hove distribution function of the sticker residues, *G*_*s*_(*r, t*). The color scheme of dark color shows earlier distribution (*t* ∼ 10*τ*), whereas light colors are the late time behavior (*t* ∼ 10^6^*τ*). All the curves are taken in the same time interval. Gaussian fits are shown with the early time (*t* ∼ 10*τ*, blue dashed line) and late time fit (*t* ∼ 10^6^*τ*, red solid line). (A-C) Chains with perfectly periodic sticker patterns with periods 10, 25, and 100. (D) 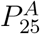 Chains with sticker periodicity 25 with shuffled stickers (E) 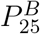 Chains with sticker periodicity 25 with alternating sticker-spacers

### Density dependent condensate architectures

The energy landscapes of stickers not only encode the final equilibrium structural and dynamical properties but also impact the time scales and percolation characteristics during the formation of condensates in the dilute biomolecular regime. We carried out simulations in the dilute regime observing a network-like fluid structure, ^31^ whereas, in the dense phase, sticky domains form percolated membrane-like structures (SI Fig. S2). In the intermediate state, where chain densities are low, spherical micelles form, and at higher densities, micelles are no longer spherical but elongated along an axis. As density increases, these elongated tube-like micelles form large branch micelles. Above a critical density, these branched micelles connect and form a single percolated porous foam-shaped membrane-like architecture.

To quantitatively characterize the shapes of the sticky clusters, we identify the clusters and calculate the gyration tensor 𝒬 _*αβ*_ of the cluster as a function of chain concentrations *ρ*. The principal components of the gyration tensor, 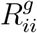 (*i* = *x, y, z*, denoted by three distinct markers), are plotted as a function of *ρ*, where for chain lengths *N*_*p*_ = 25, 50, 100 (Fig. 8 G). The vertical line separates the percolation and disperse clusters. Interestingly, all the orthogonal components of 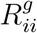 are equivalent at lower densities. However, 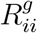 degenerates as density increases, ensuring the clusters form an elongated shape. This behavior is persistent with different chain lengths (Fig. 8 G).

**Figure 8:**
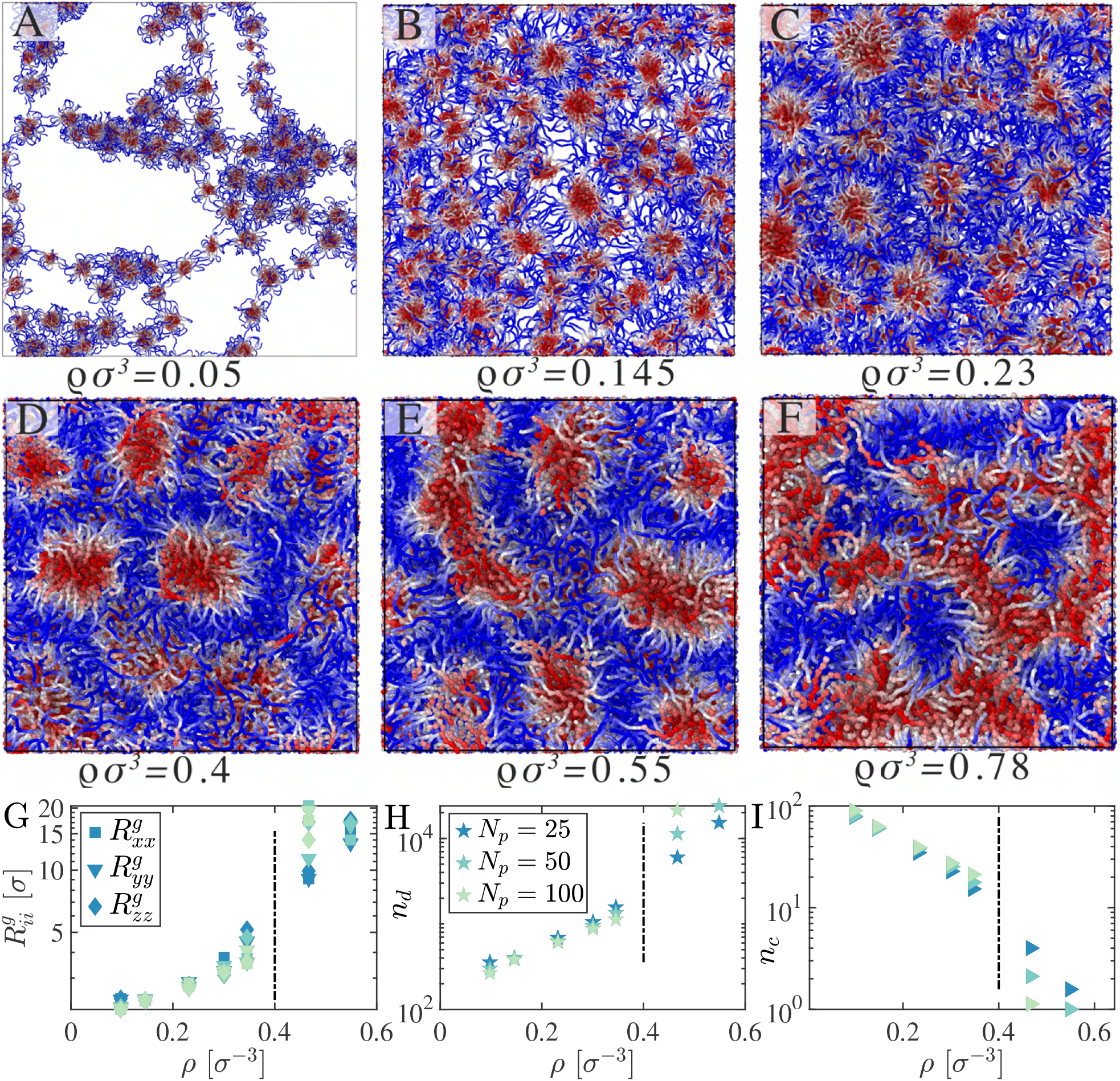
Biomolecular condensate phases of *P*_25_ at different number densities *ρ* (by varying the box volume *L*^3^) creates a range of architectures, from a network fluid structure to clusters of micelles. (G) Orthogonal radius of gyration of sticky clusters 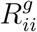, (H) average number of motifs in a single cluster, *n*_*d*_, and (I) the average number of sticky clusters in bulk, *n*_*c*_, as a function of box dimensions *ρ* are shown here. The vertical lines separate the percolated single cluster and multiple clusters in the bulk phase.

The average number of particles in each cluster *n*_*d*_ as a function of *ρ* (Fig. 8 H) for different *N*_*p*_, shows similar behavior. It is also interesting to examine the number of clusters in the system as a function of chain density, which linearly increases with density (Fig. 8 I), irrespective of the chain length. The important length scale here is the size of the cluster or the width of the membrane-like percolated surface shape, which can be quantified by identifying the anomalous peak in *S*(*q*).

To explore the morphology of clusters we use shape characteristics such as asphericity, 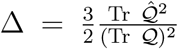and nature of asphericity, 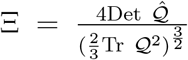 where 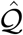 is the gyration tensor. The parameter 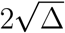, ranging from 0 to 2, characterizes cluster deviation from spherical to elongated shapes, with extreme values representing spheres and rigid rods, respectively. Whereas cos^−1^ Ξ delineates the transition between oblate and prolate shapes. We report distinct micell-like sticky cluster shapes (Fig. 9),: i) four different chain lengths *N*_*p*_ (Fig. (9 A-D)) ii) different chain densities *ρ*. Notably, the red line delineates the structural diversity within clusters between accessible and inaccessible closed shapes. We vary the density *ρ* for each chain length *N*_*p*_, indicating that spherical sticky clusters form at lower densities (*ρ* ≤ 0.2). In contrast, the sticky clusters form elongated micellar structures at higher densities (Fig. 9). The histogram of the radius of gyration *R*_*g*_ of these sticky clusters further supports this observation (Fig. 9). Under dilute conditions, equilibrium clusters are smaller, exhibit more fluid-like behavior, and are loosely connected within the network (see SI, Fig S4). Conversely, at higher densities, the cluster node size increases, and the sticky nodes are tightly connected, resulting in a more elastic network.

**Figure 9:**
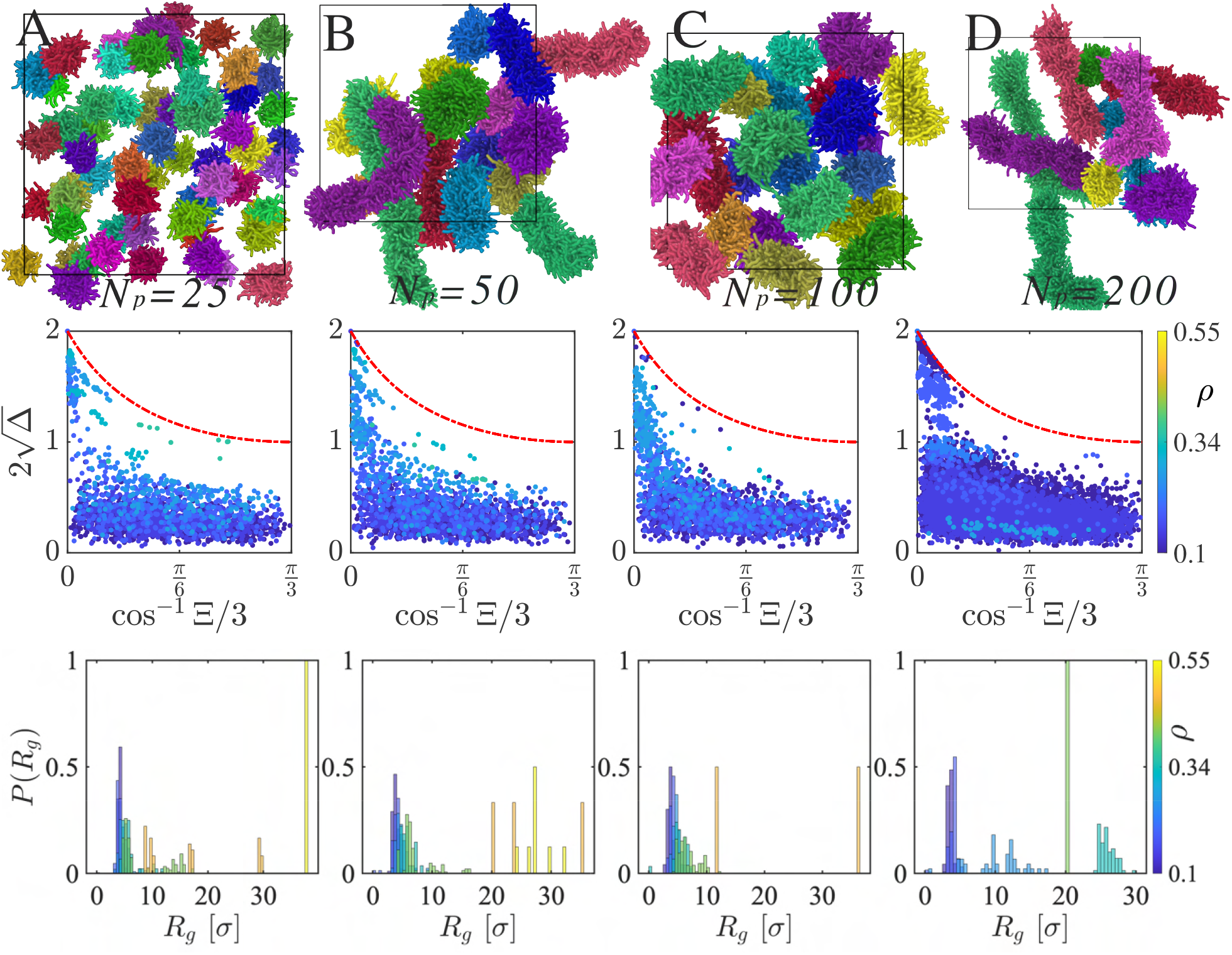
The first row displays snapshots of sticker clusters for four different chain lengths *N* : (A) 25, (B) 50, (C) 100, and (D) 200, with individual clusters depicted in various colors. In the second row, corresponding systems for each chain length illustrate how cluster shapes evolve across different density regimes, transitioning from spherical to elongated micelles. The third row presents the probability distribution of the radius of gyration, *P* (*R*_*g*_), for the sticker clusters.

## Discussion

A significant fraction of eukaryotic proteins contain low-complexity regions with unknown functions. ^2^ Mounting evidence suggests an important functional utility of LCDs in proteins for driving phase separation through associative interactions. ^29,51,52^ A paradigmatic architecture of condensate-forming proteins has two key elements: stickers, which could be either LCDs with stretches of charged/aromatic domains or foldable domains with attractive patches and spacers, which are flexible regions often enriched glycines.^1,39,40^ Decoding how sticker patterns encode the material properties of condensates promises to shed light on the dynamic behavior of condensates, which are intimately linked to their sequence patterns and cellular functions. ^4,53,54^

In this work, we employ an energy landscape framework of stickers and spacers to investigate the sticker/spacer patterns that may be evolutionarily constrained to possess both short-term material properties and long-term stability required for cellular functions. The energy landscape framework allows us to analyze how periodic and quasi-periodic sticker/spacer patterns and sticker affinities affect the equilibrium and nonequilibrium rheology of homotypic protein condensates. By continuously evolving sequence types, we highlight four critical extremes based on the periodicity of sticker and spacer motifs: i) extreme periodic motifs, fully randomized motifs, iii) randomness within the sticker regions, and iv) alternating sticker and spacer patterns.

Our simulations show that variation of a handful of energy landscape features can capture viscoelastic trends in recent experiments on prion domains.^10,19,42^ We predict how one can systematically program protein condensate viscoelasticity by altering the periodicity and strength of sticker motifs. Guided by the energy landscape framework, we continuously evolve sequences from extreme periodic to extreme random distribution of stickers and spacers. We find periodic repeats of low-complexity sticker regions display local structure and inhomogeneities in the bulk system. In contrast, random sequences show homogeneous unstructured bulk systems, which are more viscous, as demonstrated by the order of magnitude difference in viscous moduli compared to the sequences with more periodic patterns of stickers. Phase diagrams and dynamical properties also corroborate the rheological trends; in the case of periodic sticker patterns, elastic response, and viscosity get higher and become more subdiffusive with an increasing periodicity of the sticker motifs.

Structurally, sticker-sticker interactions in periodic sequence motifs lead to tangent correlation’s distinct oscillatory exponential decay. The van-Hove analysis shows that high sequence patterns with a high degree of periodicity can lead to glassy arrested systems emphasizing the slow dynamics. By varying densities of the system, we shed light on distinct architectures formed by different degrees of periodic order. We classify the shape of the sticker cluster; in low densities, a spherical micelles-like structure has been observed; however, in higher densities, we observed elongated micelles, which form branched percolated micelles at even higher concentrations.

Our study bridges a critical gap between sequence grammar and material properties of biomolecular condensates by quantitatively and qualitatively dissecting the impact of periodic sticker and spacer motifs on structure, dynamics and observable rheological quantities. These findings contribute to the fundamental understanding of the equilibrium properties of condensates but also offer potential avenues for manipulating structural behaviors in sticker spacer models.

## Simulation Methods

Molecular dynamics simulations were conducted using the LAMMPS package. The equations of motion integrated via the NVT and Langevin thermostat employing a time step of 0.001*τ*, where 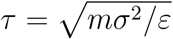, represents the derived unit of time and m denotes the mass of a bead. Langevin thermostat was used with a friction factor of 0.1*m/τ*. Chains of fixed length *N*_*p*_ = 200 were initially positioned randomly within a periodic cubic simulation box with edge lengths of *L*_*x*_ = *L*_*y*_ = *L*_*z*_ = 50*σ*. To compute the phase diagram, the simulation box was expanded along the *x*−axis by a factor of 5 (resulting in *L*_*x*_ = 250*σ*), yielding a slab of condensed phase. Phase diagram was calculated based on the density profile of the slab. The critical point was estimated via extrapolation using rectilinear diameters and the universal scaling of coexistence densities (Fig. 3 A), approaching asymptotically close to the critical point, expressed as 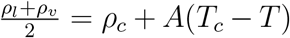 and Δ*ρ* = Δ*ρ*_0_ + (1 − *T/T*_*c*_)^*β*^, with the exponent *β* = 0.325.

For viscoelasticity calculations we equilibrated the systems inside a cubic box *L* = 50*σ*. The Green-Kubo approach have been implemented to extract viscoelastic properties. We used the multi-tau correlator method to effectively reduce noise while preserving accurate relaxation dynamics.^55,56^ From the equilibrium stress autocorrelation function of an isotropic system, one can approximate complex modulus as:

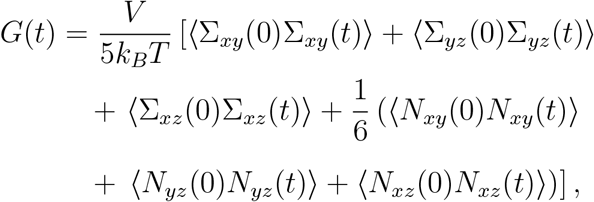

where Σ_*ij*_ represents the off-diagonal components of the stress tensor, and *N*_*ij*_ = Σ_*ii*_ − Σ_*jj*_ denotes the normal stress tensor difference. Complex viscosity, denoted as *η*, is calculated using 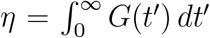, where the complex modulus *G* is defined as *G* = *G*^*′*^ + *iG*^*′′*^.^57,58^ For viscoelastic fluids, the real part of the complex modulus, *G*^*′*^, the elastic modulus, also known as the storage modulus or in-phase modulus, quantifies the solid-like response within the system. Whereas the imaginary part, *G*^*′′*^, the viscous modulus, termed the loss modulus or out-of-phase modulus, is associated with energy dissipation through viscous flow. Fourier transform of the relaxation time dependent complex modulus 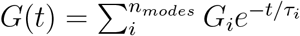, gives frequency dependent generalized Maxwell mode via 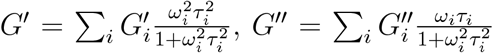 The viscosity of viscoelastic systems can be determined as 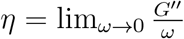.

## Acknowledgement

The authors acknowledge the support from the Research Corporation for Scientific Advancement Cottrell Scholar Award. DP also acknowledges support from the National Institutes of Health with grant no R35GM138243.

## Supporting Information Available

Additional data

## References

(1) Holehouse, A. S.; Kragelund, B. B. The molecular basis for cellular function of intrinsically disordered protein regions. Nature reviews Molecular cell biology 2024, 25, 187–211.

(2) McKnight, S. L. Protein domains of low sequence complexity—dark matter of the proteome. Genes & development 2024, 38, 205–212.

(3) Bhowmick, A.; Brookes, D. H.; Yost, S. R.; Dyson, H. J.; Forman-Kay, J. D.; Gunter, D.; Head-Gordon, M.; Hura, G. L.; Pande, V. S.; Wemmer, D. E.; others Finding our way in the dark proteome. Journal of the American Chemical Society 2016, 138, 9730–9742.

(4) Zhou, H.-X.; Kota, D.; Qin, S.; Prasad, R. Fundamental Aspects of Phase-Separated Biomolecular Condensates. Chemical Reviews 2024, 124, 8550–8595.

(5) Linsenmeier, M.; Hondele, M.; Grigolato, F.; Secchi, E.; Weis, K.; Arosio, P. Dynamic arrest and aging of biomolecular condensates are modulated by low-complexity domains, RNA and biochemical activity. Nat. Commun. 2022, 13, 3030.

(6) Niaki, A. G.; Sarkar, J.; Cai, X.; Rhine, K.; Vidaurre, V.; Guy, B.; Hurst, M.; Lee, J. C.; Koh, H. R.; Guo, L.; others Loss of dynamic RNA interaction and aberrant phase separation induced by two distinct types of ALS/FTD-linked FUS mutations. Molecular cell 2020, 77, 82–94.

(7) Michaels, T. C.; Qian, D.; Šarić, A.; Vendruscolo, M.; Linse, S.; Knowles, T. P. Amyloid formation as a protein phase transition. Nature Reviews Physics 2023, 5, 379–397.

(8) Alberti, S.; Hyman, A. A. Biomolecular condensates at the nexus of cellular stress, protein aggregation disease and ageing. Nature reviews Molecular cell biology 2021, 22, 196–213.

(9) Alshareedah, I.; Moosa, M. M.; Pham, M.; Potoyan, D. A.; Banerjee, P. R. Programmable viscoelasticity in protein-RNA condensates with disordered sticker-spacer polypeptides. Nature communications 2021, 12, 6620.

(10) Alshareedah, I.; Singh, A.; Yang, S.; Ramachandran, V.; Quinn, A.; Potoyan, D. A.; Banerjee, P. R. Determinants of viscoelasticity and flow activation energy in biomolecular condensates. Sci Adv 2024, 10, eadi6539.

(11) Sundaravadivelu Devarajan, D.; Wang, J.; Szała-Mendyk, B.; Rekhi, S.; Nikoubashman, A.; Kim, Y. C.; Mittal, J. Sequence-dependent material properties of biomolecular condensates and their relation to dilute phase conformations. Nature Communications 2024, 15, 1912.

(12) Tejedor, A. R.; Collepardo-Guevara, R.; Ramírez, J.; Espinosa, J. R. Time-dependent material properties of aging biomolecular condensates from different viscoelasticity measurements in molecular dynamics simulations. The Journal of Physical Chemistry B 2023, 127, 4441–4459.

(13) Rana, U.; Xu, K.; Narayanan, A.; Walls, M. T.; Panagiotopoulos, A. Z.; Avalos, J. L.; Brangwynne, C. P. Asymmetric oligomerization state and sequence patterning can tune multiphase condensate miscibility. Nature Chemistry 2024, 1–10.

(14) Rekhi, S.; Garcia, C. G.; Barai, M.; Rizuan, A.; Schuster, B. S.; Kiick, K. L.; Mittal, J. Expanding the molecular language of protein liquid–liquid phase separation. Nature Chemistry 2024, 1–12.

(15) Lytle, T. K.; Chang, L.-W.; Markiewicz, N.; Perry, S. L.; Sing, C. E. Designing electrostatic interactions via polyelectrolyte monomer sequence. ACS central science 2019, 5, 709–718.

(16) Choi, J.-M.; Holehouse, A. S.; Pappu, R. V. Physical principles underlying the complex biology of intracellular phase transitions. Annual review of biophysics 2020, 49, 107–133.

(17) Ginell, G. M.; Holehouse, A. S. Phase-Separated Biomolecular Condensates: Methods and Protocols; Springer, 2022; pp 95–116.

(18) Biswas, S.; Potoyan, D. A. Molecular Drivers of Aging in Biomolecular Condensates: Desolvation, Rigidification, and Sticker Lifetimes. PRX Life 2024, 2, 023011.

(19) Alshareedah, I.; Borcherds, W. M.; Cohen, S. R.; Singh, A.; Posey, A. E.; Farag, M.; Bremer, A.; Strout, G. W.; Tomares, D. T.; Pappu, R. V.; others Sequence-specific interactions determine viscoelasticity and ageing dynamics of protein condensates. Nature Physics 2024, 1–10.

(20) Le, N. T. K.; Park, E.; Kim, H.; Park, J.; Kang, K. Viscosity Regulation of Chemically Simple Condensates. Biomacromolecules 0, 0, null.

(21) Onuchic, J. N.; Luthey-Schulten, Z.; Wolynes, P. G. Theory of protein folding: the energy landscape perspective. Annual review of physical chemistry 1997, 48, 545–600.

(22) Bryngelson, J. D.; Onuchic, J. N.; Socci, N. D.; Wolynes, P. G. Funnels, pathways, and the energy landscape of protein folding: a synthesis. Proteins: Structure, Function, and Bioinformatics 1995, 21, 167–195.

(23) Sali, A.; Shakhnovich, E.; Karplus, M. How does a protein fold? nature 1994, 369, 248–251.

(24) Guo, H.-B.; Perminov, A.; Bekele, S.; Kedziora, G.; Farajollahi, S.; Varaljay, V.; Hinkle, K.; Molinero, V.; Meister, K.; Hung, C.; others AlphaFold2 models indicate that protein sequence determines both structure and dynamics. Scientific Reports 2022, 12, 10696.

(25) Bowman, G. R. AlphaFold and Protein Folding: Not Dead Yet! The Frontier Is Conformational Ensembles. Annual Review of Biomedical Data Science 2024, 7, 11.

(26) Roney, J. P.; Ovchinnikov, S. State-of-the-Art Estimation of Protein Model Accuracy Using AlphaFold. Phys. Rev. Lett. 2022, 129, 238101.

(27) Collepardo-Guevara, R.; Joseph, J. A.; Wales, D. J.; others Energy landscapes and heat capacity signatures for peptides correlate with phase separation propensity. QRB discovery 2023, 4, e7.

(28) Chen, F.; Jacobs, W. M. Emergence of multiphase condensates from a limited set of chemical building blocks. Journal of Chemical Theory and Computation 2023,

(29) Pappu, R. V.; Cohen, S. R.; Dar, F.; Farag, M.; Kar, M. Phase transitions of associative biomacromolecules. Chemical Reviews 2023, 123, 8945–8987.

(30) Ramachandran, V.; Potoyan, D. A. Energy landscapes of homopolymeric RNAs revealed by deep unsupervised learning. Biophysical Journal 2024, 123, 1152–1163.

(31) Dar, F.; Cohen, S. R.; Mitrea, D. M.; Phillips, A. H.; Nagy, G.; Leite, W. C.; Stanley, C. B.; Choi, J.-M.; Kriwacki, R. W.; Pappu, R. V. Biomolecular condensates form spatially inhomogeneous network fluids. Nat. Commun. 2024, 15, 3413.

(32) Holehouse, A. S.; Pappu, R. V. Functional implications of intracellular phase transitions. Biochemistry 2018, 57, 2415–2423.

(33) Bracha, D.; Walls, M. T.; Wei, M.-T.; Zhu, L.; Kurian, M.; Avalos, J. L.; Toettcher, J. E.; Brangwynne, C. P. Mapping local and global liquid phase behavior in living cells using photo-oligomerizable seeds. Cell 2018, 175, 1467–1480.

(34) Watanabe, F.; Akimoto, T.; Best, R. B.; Lindorff-Larsen, K.; Metzler, R.; Yamamoto, E. Diffusion of intrinsically disordered proteins within viscoelastic membraneless droplets. arXiv 2024,

(35) Jawerth, L.; Fischer-Friedrich, E.; Saha, S.; Wang, J.; Franzmann, T.; Zhang, X.; Sachweh, J.; Ruer, M.; Ijavi, M.; Saha, S.; others Protein condensates as aging Maxwell fluids. Science 2020, 370, 1317–1323.

(36) Cohen, S. R.; Banerjee, P. R.; Pappu, R. V. Direct computations of viscoelastic moduli of biomolecular condensates. bioRxiv 2024, 2024–06.

(37) Zhang, Y.; Prasad, R.; Su, S.; Lee, D.; Zhou, H.-X. Amino Acid-Dependent Material Properties of Tetrapeptide Condensates. bioRxiv 2024,

(38) Galvanetto, N.; Ivanović, M. T.; Del Grosso, S. A.; Chowdhury, A.; Sottini, A.; Nettels, D.; Best, R. B.; Schuler, B. Mesoscale properties of biomolecular condensates emerging from protein chain dynamics. arXiv preprint 2024,

(39) Strader, R. L.; Shmidov, Y.; Chilkoti, A. Encoding Structure in Intrinsically Disordered Protein Biomaterials. Accounts of Chemical Research 2024, 57, 302–311.

(40) Dzuricky, M.; Rogers, B. A.; Shahid, A.; Cremer, P. S.; Chilkoti, A. De novo engineering of intracellular condensates using artificial disordered proteins. Nature chemistry 2020, 12, 814–825.

(41) Martins, M. L.; Zhao, X.; Demchuk, Z.; Luo, J.; Carden, G. P.; Toleutay, G.; Sokolov, A. P. Viscoelasticity of polymers with dynamic covalent bonds: concepts and misconceptions. Macromolecules 2023, 56, 8688–8696.

(42) Wu, X.; Sun, Y.; Yu, J.; Miserez, A. Tuning the viscoelastic properties of peptide coacervates by single amino acid mutations and salt kosmotropicity. Communications Chemistry 2024, 7, 5.

(43) Cagiada, M.; Bottaro, S.; Lindemose, S.; Schenstrøm, S. M.; Stein, A.; Hartmann-Petersen, R.; Lindorff-Larsen, K. Discovering functionally important sites in proteins. Nature communications 2023, 14, 4175.

(44) Di Pierro, M.; Potoyan, D. A.; Wolynes, P. G.; Onuchic, J. N. Anomalous diffusion, spatial coherence, and viscoelasticity from the energy landscape of human chromosomes. Proceedings of the National Academy of Sciences 2018, 115, 7753–7758.

(45) Liu, L.; Shi, G.; Thirumalai, D.; Hyeon, C. Chain organization of human interphase chromosome determines the spatiotemporal dynamics of chromatin loci. PLoS computational biology 2018, 14, e1006617.

(46) Guenza, M. G. Anomalous Dynamics in Macromolecular Liquids. Polymers 2022, 14, 856.

(47) Van Hove, L. Correlations in Space and Time and Born Approximation Scattering in Systems of Interacting Particles. Phys. Rev. 1954, 95, 249–262.

(48) Hopkins, P.; Fortini, A.; Archer, A. J.; Schmidt, M. The van Hove distribution function for Brownian hard spheres: Dynamical test particle theory and computer simulations for bulk dynamics. The Journal of Chemical Physics 2010, 133, 224505.

(49) Chaudhuri, P.; Berthier, L.; Kob, W. Universal Nature of Particle Displacements close to Glass and Jamming Transitions. Phys. Rev. Lett. 2007, 99, 060604.

(50) Weeks, E. R.; Crocker, J. C.; Levitt, A. C.; Schofield, A.; Weitz, D. A. Threedimensional direct imaging of structural relaxation near the colloidal glass transition. Science 2000, 287, 627–631.

(51) Hardenberg, M.; Horvath, A.; Ambrus, V.; Fuxreiter, M.; Vendruscolo, M. Widespread occurrence of the droplet state of proteins in the human proteome. Proceedings of the National Academy of Sciences 2020, 117, 33254–33262.

(52) Li, P.; Chen, P.; Qi, F.; Shi, J.; Zhu, W.; Li, J.; Zhang, P.; Xie, H.; Li, L.; Lei, M.; others High-throughput and proteome-wide discovery of endogenous biomolecular condensates. Nature Chemistry 2024, 1–12.

(53) Zhou, H.-X. Viscoelasticity of biomolecular condensates conforms to the Jeffreys model. The Journal of Chemical Physics 2021, 154, 041103.

(54) Guenza, M. G. Dynamics of protein droplets revealed by bridging multiple scales. Nature 2023, 619, 700–701.

(55) Ramírez, J.; Sukumaran, S. K.; Vorselaars, B.; Likhtman, A. E. Efficient on the fly calculation of time correlation functions in computer simulations. The Journal of Chemical Physics 2010, 133, 154103.

(56) Thompson, A. P.; Aktulga, H. M.; Berger, R.; Bolintineanu, D. S.; Brown, W. M.; Crozier, P. S.; in‘t Veld, P. J.; Kohlmeyer, A.; Moore, S. G.; Nguyen, T. D.; Shan, R.; Stevens, M. J.; Tranchida, J.; Trott, C.; Plimpton, S. J. LAMMPS - a flexible simulation tool for particle-based materials modeling at the atomic, meso, and continuum scales. Comp. Phys. Comm. 2022, 271, 108171.

(57) Ferry, J. D. Viscoelastic properties of polymers; John Wiley & Sons, 1980.

(58) Shaw, M. T.; MacKnight, W. J. Introduction to polymer viscoelasticity; John Wiley & Sons, 2018.

