## Supporting Information for "Decoding Biomolecular Condensate Dynamics: An Energy Landscape Approach"

### A. Contact map

The spatial correlation can be inferred from the Contact map map, which leads to probing the folded domain information between residues within the sticker-spacer model. Which would help understand and visualize AFELAS chain structures and understand their interactions. Typically, the contact maps depict pairwise interactions between residues or atoms within a protein structure, offering insights into critical interactions stabilizing different conformations and controlling protein functions.

To calculate Contact maps, structural models are obtained from equilibrium trajectory files, and residue coordinates are extracted. Contact maps are then generated for each periodically or its randomly distributed sticker-spacer chains, considering residues to be in contact if the distance between any two $i$ and $j$ the residue satisfies $r < r_c = 2\sigma$. This approach provides a means to analyze and compare protein structures, particularly beneficial for understanding the impact of mutations on protein function and conformational flexibility.

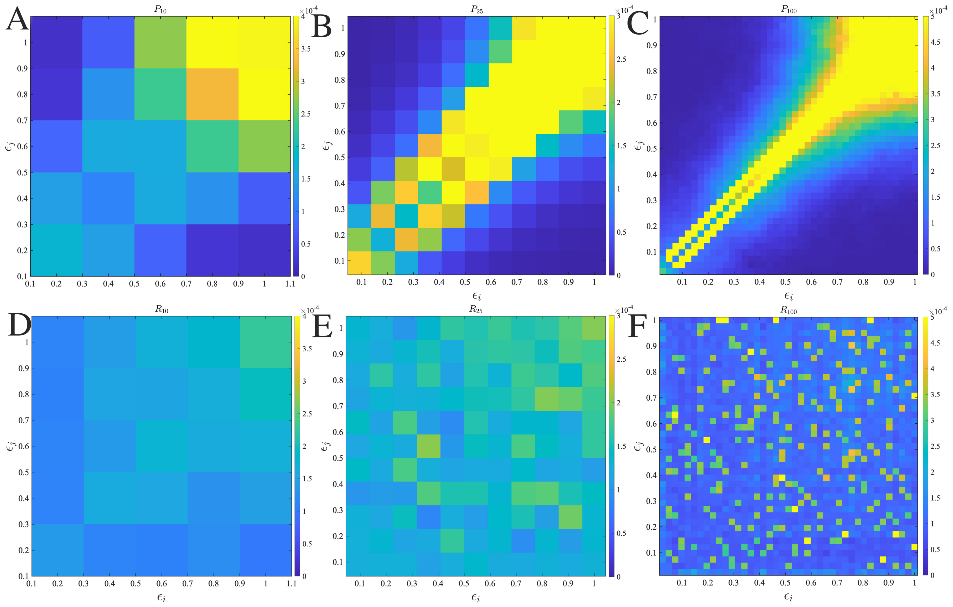


FIG. S1. Contact maps illustrate the spatial proximity $r_c < 2\sigma$ between motifs $i$ and $j$ within a biomolecular condensate in equilibrium for PDSSM (a) $P_{10}$ (b) $P_{25}$ (c) $P_{100}$ and RDSSM (d) $R_{10}$ (e) $R_{25}$ (f) $R_{100}$. In the case of PDSSM, sticker motifs have formed a cluster, which is highest in sticker residues. While there is a lack of consistent contact between spacers. Conversely, RDSSM exhibits no consistent contact regions capable of forming a sticker domain.

As shown in Fig. (S1), we calculate the Contact map of AFELAS chins in the bulk phase, periodically distributed sticker-spacer motifs (A) $P_{10}$ (B) $P_{25}$ (C) $P_{100}$. It is clear from the figures that stickers that have higher interaction strengths $\varepsilon_i$ try to form contacts that are stable and form folded domains, which makes these contacts even stronger. For example, $P_{100}$ has more stable contacts compare to $P_{25}$ and $P_{10}$. However, while the residues are randomly

*
†

shuffled for randomly distributed sticker-spacer motifs (D) $R_{10}$ (E) $R_{25}$ (F) $R_{100}$, contacts are spatially uncorrelated and not stable.

#### B. Network Fluid

Proteins with associative properties exhibit distinct molecular traits, such as oligomerization domains, ligand binding domains, and intrinsically disordered regions. The coupling of associative and segregative phase transitions is driven by these molecular features. The formation of higher-order complexes eventually results in the segregation into dilute and dense phases. Percolation, like bond percolation or physical gelation, facilitates the formation of network-like structures through reversible physical crosslinks. These networks, influenced by molecular architectures and crosslinking, contribute to the viscoelastic properties of condensates. In this work, we report that the condensed phases show different architectures of periodically distributed residue chains with varying densities $\rho$. In very dilute solutions, stickers among single chains individually collapse and form dispersed globular chains in the system. In a dense phase, the sticker cluster is percolated in the bulk. Whereas, in intermediate lower densities, sticker motifs form smaller monodisperse clusters, which are connected via spacers for longer chains. How these sticker clusters are connected in the bulk phase can be quantified using a graph neural network. the graph network in Fig. (S2), we show how sticky clusters are connected, with different densities (by varying box dimension $L_b$) and chain lengths $N_p$ of $P_{25}$ AFELAS chians. It is clear from the chart that sticky cluster nodes are not connected for shorter chains $N_p = 25$ (matches the residue periodicity), which is independent of system density. For chains $N_p = 50, 100, 200$, we see distinct nodes that are connected for lower densities. As chain length increases number of nodes increases. However, connectivity is stronger at some intermediate densities , *i.e.*, $L_b = 60$; however, for even higher densities,

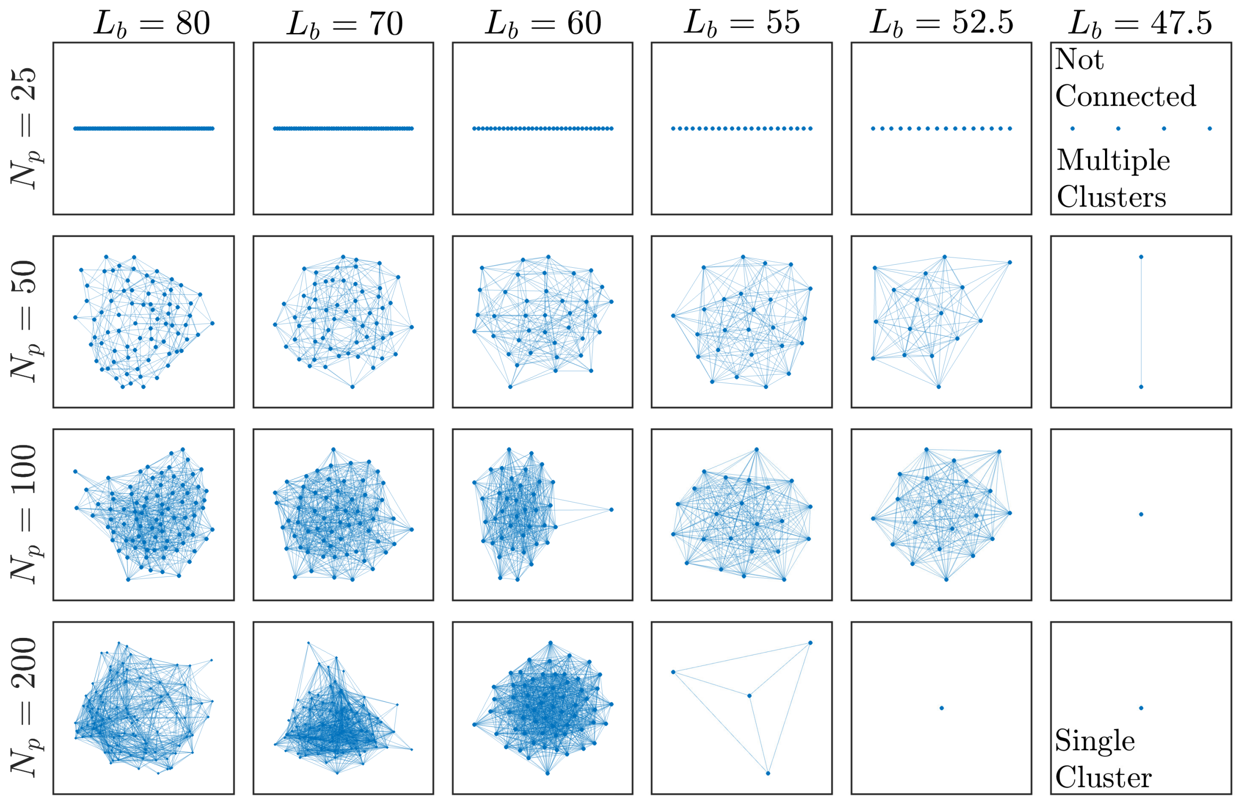


FIG. S2. Connected network of sticky clusters for $P_{25}$ AFELAS chains are denoted by nodes, and the connection between node stickers are connected via edge lines. For denser systems, the sticker cluster forms one membrane-like percolated surface, which is denoted by a single cluster. In the phase space of $N_p$ and $L_b$ or the density, $N_p = 25$ shorter chains do not form a connected network.

there are few nodes that are connected or a single cluster that is percolated through the system.

The figure in Fig. (S3) illustrates the radius of gyration ($R_g$) for the sticker clusters of AFELAS $P_{25}$ chains across various densities. In the dilute regime, depicted in Fig. S3(A), clusters formed by stickers appear smaller initially, gradually enlarging as more chains join within a single cluster. This phenomenon results in tightly interconnected networks, influencing the viscoelastic properties. With increasing density, the size of sticker clusters expands, leading to a shift in the mean of the $R_g$ distribution towards higher values.

Furthermore, in the dense phase, our observations indicate the breakdown of spherical symmetry, with the formation of elongated micelles or even branched micelles at higher densities. Beyond a certain cutoff radius, all stickers are interconnected through a percolated structure, resulting in a delta function value for the radius of gyration, as depicted in Fig. S3(F).

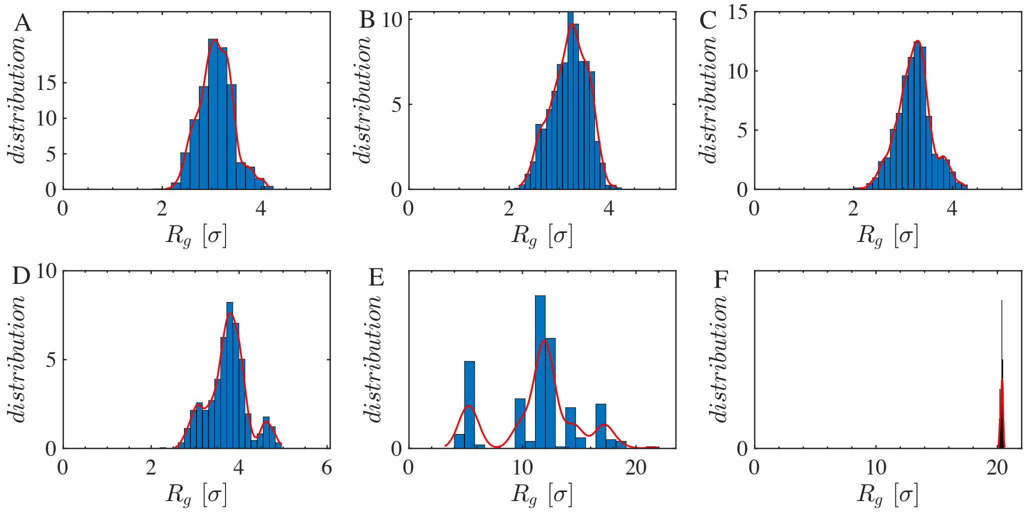


FIG. S3. Distribution of Radius of gyration $P(R_g)$ of the cluster as a function of the volume fraction of the biomolecular systems AFELAS $P_{25}$ chains with $N_p = 200$.

#### C. Contact time

Given the pronounced sensitivity of interaction strength among interactive residues within the dense phases of patterned residue sequences to their specific arrangements, we probe into the dynamics of contact formation. This approach enables direct comparison with the mechanical properties of sticker-spacer residue condensates. We calculate of the intermittent contact time autocorrelation function $C(t)$, expressed as:

$$C(t) = \frac{\langle H(t_0)H(t+t_0)\rangle}{\langle H(t_0)H(t_0)\rangle},$$

where $H$ is a step function defined to be one if a pair of interactive residues $i$ and $j$ on two different residues are within a cutoff radius (smaller than $r_c$, the average diameter of residues $i$ and $j$) at both initial time $t = 0$ and at time $t$, and 0 otherwise. This autocorrelation function captures the instances when contacts between residues are broken and subsequently reformed, thus reflecting the dynamic stability of these interactions within the condensate.

Here $C(t)$, time-averaged over all interacting residue pairs, indicated multiple decay behaviors with varying degrees of residue segregation in stickers and spacers with varying sequence pattering. To further analyze these dynamics, we computed the intermittent contact lifetime $\tau$ by integrating $C(t)$ until it decreased to a value of a very long time.

The autocorrelation function, $C(t)$, not only reflects the dynamics of contact formation but also provides indirect insights into the viscoelastic properties of protein condensates. At short timescales, $C(t)$ tends to decay rapidly, indicative of fluid-like behavior characterized by the frequent breaking and reforming of contacts. In the case of periodic residues $P_{25}$ and $P_{25}^A$, longer timescale behavior $C(t)$ suggests a transition to a more gel-like state, marked by sustained interactions that enhance the overall viscosity and structural rigidity of the condensate. This transition is primarily due to the arrest of sticker-sticker interactions. As shown in Fig. (S4 A) $C(t)$ plotted as a function of time for different temperatures and the different pattering of stickers and spacers, which plays a crucial role in determining the material properties of the condensates, effectively bridging molecular interactions with macroscopic

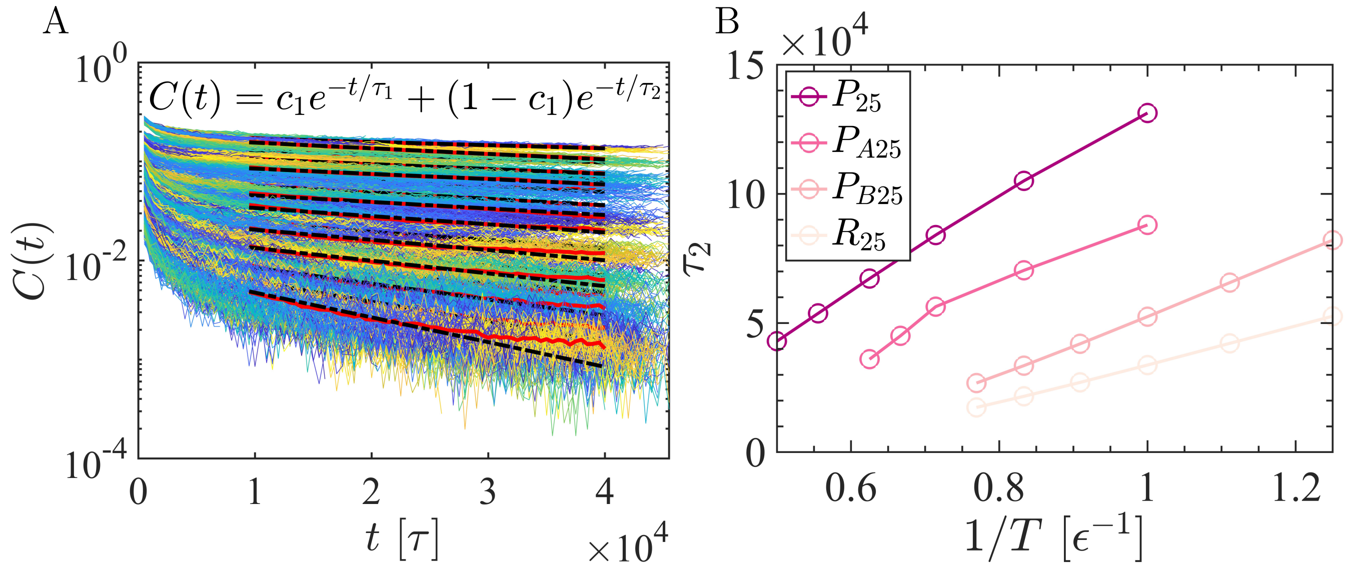


FIG. S4. (a) The contact time auto-correlation function $C(t)$, as a function of time for different sticker-spacer arrangements for $P_{25}$, $P_{A25}$, $P_{B25}$, and $R_{25}$ is shown here. (b) The longer correlation time $\tau_2$ is plotted as a function of inverse temperature $1/T$. For periodic chains, $P_{25}$ has a much stronger correlation, which even increases at lower temperatures. The correlation time $\tau_2$ decreases systematically for $P_{A25}$ and $P_{B25}$. The correlation time is lowest for randomly arranged residues $R_{25}$.

physical properties. When stickers are periodically arranged, $C(t)$ exhibits long correlation times. Nonetheless, an increase in temperature $T$ accelerates the decay of long-time correlations, indicating a reduction in the viscoelastic stabilization contributed by sticker interactions as shown in Fig. (S4 B).
